# Long-read sequencing to interrogate strain-level variation among adherent-invasive *Escherichia coli* isolated from human intestinal tissue

**DOI:** 10.1101/2020.03.10.985440

**Authors:** Jeremy Wang, Rachel Bleich, Sandra Zarmer, Shiying Zhang, Belgin Dogan, Kenneth W Simpson, Janelle Arthur

## Abstract

Adherent-invasive *Escherichia coli* (AIEC) is a pathovar linked to inflammatory bowel diseases (IBD), especially Crohn’s disease, and colorectal cancer. AIEC are genetically diverse, and in the absence of a universal molecular signature, are defined by *in vitro* functional attributes. The relative ability of difference AIEC strains to colonize, persist, and induce inflammation in an IBD-susceptible host is unresolved. To evaluate strain-level variation among tissue-associated *E. coli* in the intestines, we develop a long-read sequencing approach to identify AIEC by strain that excludes host DNA. We use this approach to distinguish genetically similar strains and assess their fitness in colonizing the intestine. Here we have assembled complete genomes using long-read nanopore sequencing for a model AIEC strain, NC101, and seven strains isolated from the intestinal mucosa of Crohn’s disease and non-Crohn’s tissues. We show these strains can colonize the intestine of IBD susceptible mice and induce inflammatory cytokines from cultured macrophages. We demonstrate that these strains can be quantified and distinguished in the presence of 99.5% mammalian DNA and from within a fecal population. Analysis of global genomic structure and specific sequence variation within the ribosomal RNA operon provides a framework for efficiently tracking strain-level variation of closely-related *E. coli* and likely other commensal/pathogenic bacteria impacting intestinal inflammation in experimental settings and IBD patients.

## Introduction

Crohn’s disease (CD) is a type of Inflammatory bowel disease (IBD), a chronic inflammatory condition that is the result of an inappropriate immune response towards elements of the intestinal microbiota [1]. Due to chronic inflammation and exposure to pro-carcinogenic microbes, CD patients are at a greater risk of developing colorectal cancer (CRC) [2–4]. CD patients harbor microbiomes that differ from non-CD individuals in both composition and function, with reduced diversity and an increase of <u>a</u>dherent/<u>i</u>nvasive *<u>E</u>. <u>c</u>oli* (AIEC) [5–11]. AIEC are closely associated with the intestinal mucosa, termed the mucosal niche, and induce inflammation in murine models of IBD, including the well-established, inflammation-susceptibe *Il10*^-/-^ mouse model [3, 12–14]. AIEC strains lack genes associated with virulence in diarrheagenic *E. coli* and are defined through the *in vitro* ability to adhere to and invade epithelial cells and survive in macrophages [5, 7, 15]. However, as AIEC are genetically diverse, there is no universal genomic feature that distinguishes the pathotype [16], limiting our ability to detect these pro-inflammatory strains from among patient microbiota.

*Escherichia coli* make up a diverse species that is known to variously have beneficial, commensal, or pathogenic impact in the mammalian gut. However, most commonly used methods for characterizing the composition of gut microbiota from fecal or tissue samples, notably sequencing of selected variable regions of the 16S rRNA operon, do not differentiate among these closely-related but functionally divergent types. Emerging single-molecule (third-generation/long-read) sequencing technologies including Pacific Biosciences (“PacBio”) and Oxford Nanopore Technologies (“nanopore”) produce reads dozens to hundreds of kilobases in length, often without prior amplification. Recent work [17–20] has demonstrated the utility of long reads for sequencing the entire 16S gene or entire rRNA operon, with corresponding increase in taxonomic resolution by capturing more variable sequence, including all nine variable regions of 16S, the internal transcribed spacers (ITS), and 23S. Traditional metataxonomic analysis using next-generation sequencing of one or two adjacent hypervariable regions of the 16S operon are unable to discriminate among closely-related species and genera [21], much less capture strain-level variation. We propose a full-length rRNA sequencing approach – enabled by longer reads produced by nanopore sequencing – that is capable of discriminating among closely-related strains of *E. coli*. To support the extension of these approaches to characterize strain-level variation of tissue-associated (mucosa-associated) microbiota contributing to IBD and in models of experimental colitis in mice, with a particular focus on mechanistic studies of AIEC, we produced complete genome assemblies for eight AIEC and non-AIEC *E. coli*, describe the genomic variation among these strains, particularly within the rRNA operon, and demonstrate the accurate identification of these strains in mixed *in vitro* and *in vivo* microbiota.

## Methods

### DNA extraction and nanopore sequencing

We extracted ultra-high-molecular-weight genomic DNA from liquid *E. coli* cultures using a modified phenol:chloroform protocol [22]. Briefly, 1-1.5 ml of stationary-phase liquid culture was pelleted and resuspended in TE/NaCl/Triton buffer. Cells were lysed by addition of SDS to a final concentration of 2% and Proteinase K to a final concentration of 300 mg/ml and incubated at 55°C for 30 minutes. DNA was purified twice by addition of 1x volume phenol:chloroform and phase separation. DNA was precipitated by addition of 0.1x volume sodium acetate and 2x volume isopropanol. Ultra-high-molecular weight DNA was “hooked” out with a melted glass capillary and washed in 70% ethanol, dried, and resuspended in nuclease-free water (NFW) by incubation overnight at 4°C. Multiplexed sequencing libraries were prepared for nanopore sequencing using Oxford Nanopore’s Ligation Sequencing Kit (LSK109) and Barcoding Expansion (NBD104) per the manufacturer’s recommended protocol with the following modifications [23]. Instead of SPRI bead cleanup after each ligation reaction, we added 1 volume of “clumping buffer” (9% PEG 8000, 1 M NaCl, 10 mM Tris), incubated at room temperature for 30 mins, pelleted by centrifugation at 13K rpm, washed twice with 70% ethanol, then allowed to dry and resuspended in NFW by incubating at 4°C overnight. This “bead-free” cleanup both avoids shearing and loss of high-molecular-weight DNA and selects for larger fragments in the PEG precipitations. Multiplexed samples were sequenced on R9.4.1 flow cells on either MinION or GridION with real-time basecalling using Guppy v3.2.2 in high-accuracy mode. The distribution of read lengths is shown in Figure 1.

**Fig 1.**
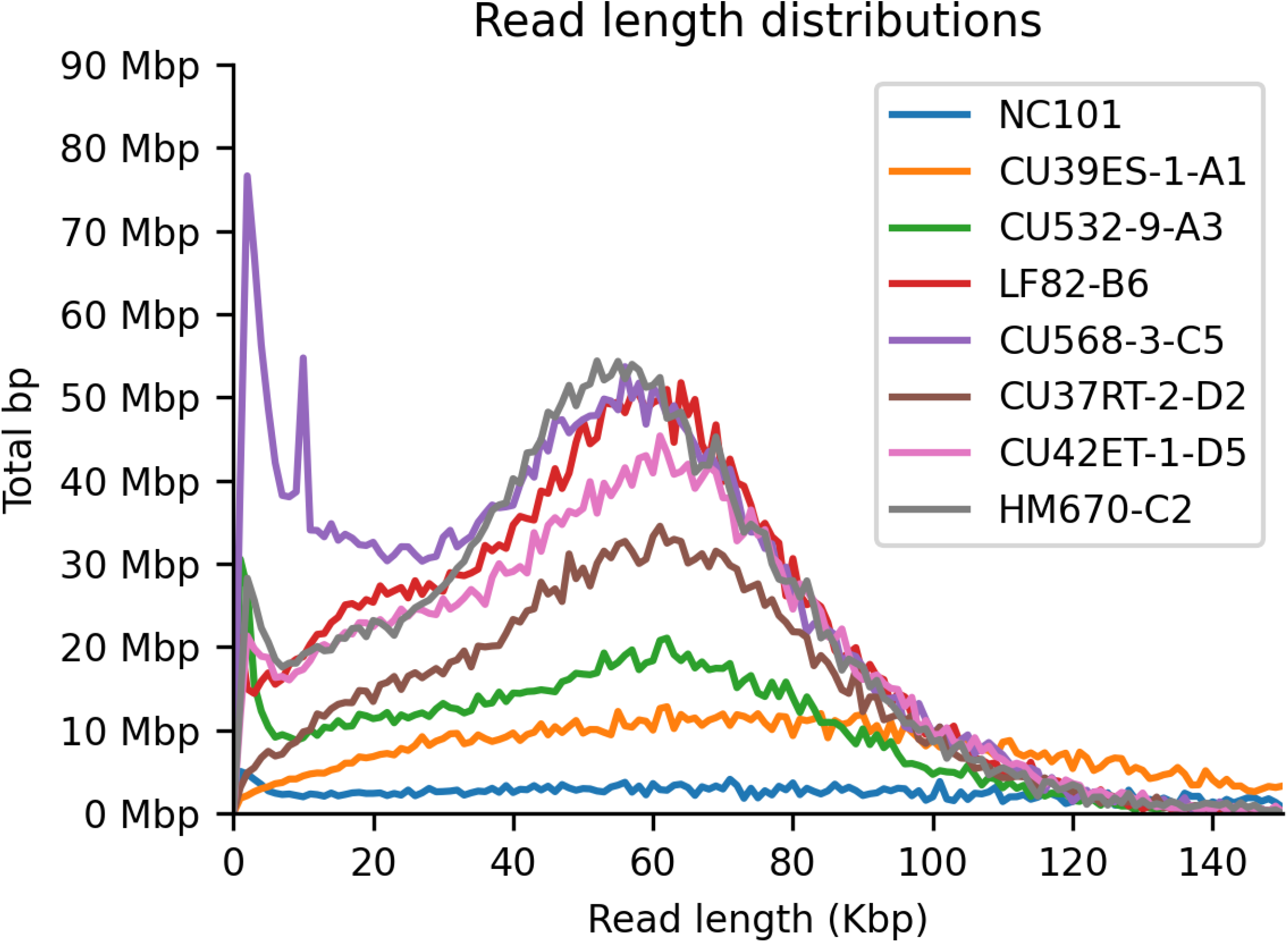
Read length distribution across samples using high-molecular weight extraction and bead-free library prep.

### Illumina sequencing

Strains NC101 and HM670 were sequenced by the Microbial Whole Genome Sequencing Center (www.migscenter.com) on the NextSeq 2000 platform, yielding paired end reads (2×151bp) delivered to us as fastq files. Strains denoted “CU” were Illumina whole-genome sequenced by Enterome (Paris, Ile-de-France, France).

### Genome assembly

Basecalled reads for each sample were assembled using Miniasm v0.2 [24], followed by four rounds of polishing with Racon v1.3.1 [25], and Medaka v0.10.0 [26]. Each assembly consisted of one contig representing the full-length genome. Assembled genomes were normalized to start at the origin of replication and re-polished across the previous breakpoint. Assemblies were further polished with Illumina sequencing data using Pilon v1.22-1 [27]. Annotation was performed using the NCBI Prokaryotic Genome Annotation Pipeline [PGAP release 2020-02-06.build4373; 28]. Predicted genes were subsequently assigned putative function by aligning to the RefSeq non-redundant protein database with Diamond v0.9.22 [29] allowing for frame-shift sensitive alignment (diamond blastp --more-sensitive --frameshift 15). Sequence data and assembled genomes have been made publicly available under NCBI BioProject ID PRJNA759208.

### Serotype, MLST, virulence, and antibiotic resistance genes

Polished assemblies were aligned against EcOH database [30] for serotyping, multi-locus sequence typing (MLST), and VirulenceFinder databases [31], and NCBI antimicrobial resistance database using Minimap2 [32] with parameter ‘-cx asm5’ to allow for indel-sensitive alignment appropriate for hybrid-assembled genomes [33]. For serotype and MLST, the single best-aligning type gene (minimum edit distance) is given as the predicted type. Predicted serotypes, MLSTs, AMR, and virulence genes were further validated using corresponding web-based tools provided by the Center for Genomic Epidemiology ([31], http://www.genomicepidemiology.org/services/), which showed complete agreement.

### rRNA operon analysis

All copies of the rRNA operon in each assembly were identified by aligning to the *E. coli* K12 reference rRNA using Minimap2 [32]. Variants relative to the K12 reference were identified for each assembled rRNA copy across all assemblies based on the alignments. Total polymorphism distance was computed between every pair of alignments across each region of the rRNA operon (16S, ITS, 23S).

### Bacterial Growth Conditions

Bacteria were grown in Luria broth (LB) at 37°C and 250 rpm unless otherwise indicated.

### Animal care

Germ-free mice were reared in the National Gnotobiotic Rodent Resource Center at UNC Chapel Hill. All animal experiments and procedures were approved by UNC’s Institutional Animal Care and Use Committee (IACUC).

### Murine stool samples

To generate an in vivo gut microbial community containing the seven *E. coli* strains of interest and endogenous mouse microbiota, we inoculated mice as follows. Three (2m/1f) interleukin-10-deficient (*Il10*^-/-^) mice (129Sv/Ev background) were reared germ-free to adulthood (8-10) weeks and colonzied by oral gavage with an even mixture of a total of 10^7^ CFU clinical *E. coli* strains isolated from the intestinal tissue of Crohn’s disease and non-Crohn’s disease patients [7]. All strains have been classified as AIEC and non-AIEC using standard *in vitro* assays to evaluate adhesion/invasion to Caco2 epithelial cells and uptake/survival in J774 macrophages [5,7]. To validate our results by tracking individual strains with PCR, each strain was marked with an antibiotic resistance cassette and molecular barcode inserted into a neutral chromosomal region using Tn7 transposon insertion [34–35]. During experimentation, these strains were referenced by their blinded laboratory designation and barcode: JA0018/A1, JA0019/A3, JA0022/B6, JA0036/C5, JA0044/D2, JA0048/D5, and JA0091/C2. The identity of these strains is described in Table 1. Five strains were originally isolated by KW Simpson at Cornell University as the strains CU39ES-1, CU532-9, CU568-3, CU37RT-2, CU42ET-1; HM670 was isolated previously [6] and gifted by Barry Campbell from Liverpool University; and LF82 is the prototypic AIEC strain gifted to KW Simpson by Arlette Darfeuille-Michaud [5].

**Table 1:**
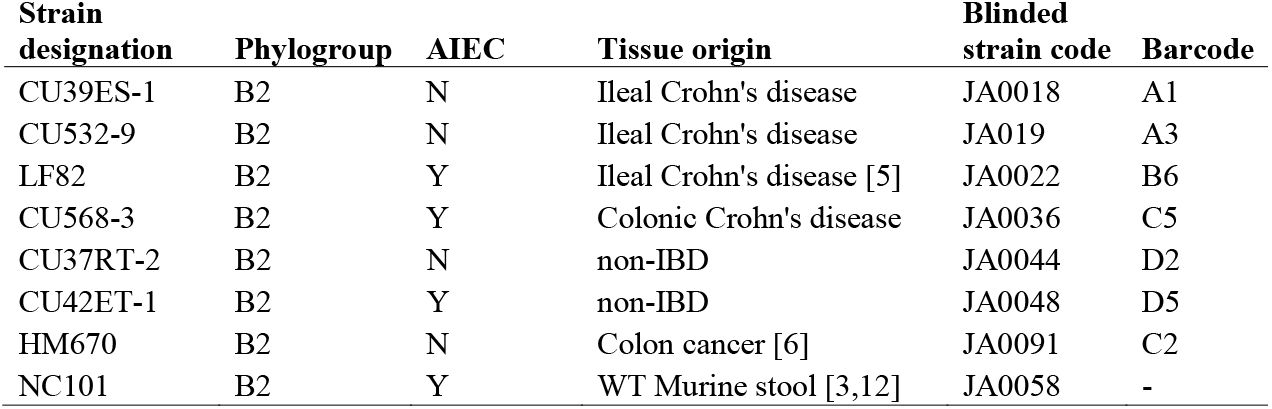
Strains used in study.

After colonization, mice were maintained in specific pathogen free (SPF) housing, where they acquired a simplified mouse microbiota. After 2 weeks of colonization, we gave kanamycin water *ad libetum* for 2 weeks to suppress this microbiota and ensure that all strains could persist to some extent. Mice were sacrificed 6 weeks later (by CO_2_ asphyxiation), for a total of 10 weeks colonization. A stool sample was removed from the lumen of the distal colon for our analysis. DNA was extracted and purified as described in [3,36].

### In vivo colonization and persistence studies

To demonstrate that the seven human-derived strains and murine *E. coli* NC101 could colonize germ-free *Il10*^-/-^ mice throughout the gastrointestinal tract, we singly housed 1 male and 1 female mouse per *E. coli* strain, for a total of 16 cages. Mice were gavaged with 10^8^ CFU of a single strain and moved to SPF housing. After 1 week, each mouse received a fecal transplant (prepared anaerobically from a pool of 7 C57BL/6 WT mice) via gavage to provide niche competition. Stool samples were collected almost daily and CFUs were quantified on kanamycin plates to monitor fecal *E. coli* colonization. Mice were sacrificed by CO2 asphyxiation after 5 weeks, and the following tissues were harvested and CFUs quantified on kanamycin plates to measure viable colonizing bacteria: ileal tissue, cecal content, colon content, colon tissue and colon mucus layer. *E. coli* CFUs were also quantified from stomach content, duodenal content, and jejunal content, but only 1-3 samples of each tissue had detectable bacterial growth of more than 10^2^ CFUs/10 mg.

To demonstrate that consistent colonizing strain, CU42ET-1-D5, and inconsistent colonizing strain, HM670-C2, colonized germ-free *Il10*^-/-^ mice in a similar manner across a larger cohort, we housed 7 (3M/4F) or 8 (4M/4F) mice, respectively, in two cages for each cohort. Mice were gavaged with 10^8^ CFU of a single strain and moved to SPF housing. After 1 week, each mouse received the same fecal transplant as used previously (prepared anaerobically from a pool of 7 C57BL/6 WT mice that were *Helicobacter spp*. free) via gavage to provide niche competition. After 3 weeks of colonization (2 weeks post-FMT), we administered kanamycin water *ad libetum* for 2 weeks to suppress this microbiota and ensure that all strains could persist to some extent. Mice were sacrificed 6 weeks later (by CO_2_ asphyxiation), for a total of 10 weeks colonization. The following tissues were harvested and CFUs quantified on kanamycin plates to measure viable colonizing bacteria: colon content, colon tissue and colon mucus layer.

### In vitro co-culture assays to measure pro-inflammatory cytokine production

To assess the inflammatory capacity of the clinical strains, J774 macrophages grown in DMEM complete media (supplemented with 10% heat-inactivated FBS and 100 U/ml penicillin-100 mg/ml streptomycin) were transferred to 12-well plates at 5×10^5^ cells/well. Cells were grown overnight at 5% CO_2_ at 37°C, and washed twice with PBS before adding fresh culture media without antibiotics. Cells were infected at an MOI of 1 for 4 hours before removing the bacteria, washing with PBS, and adding fresh media with 100 μg/mL gentamycin. Cells were grown another 20 hours at which point the supernatants were collected for ELISA and the cells were collected for qPCR analysis.

### ELISA

Supernatants from stimulated J774 macrophages (above) were analyzed for cytokine IL12p40 production through ELISA following manufacturer’s protocol (BD Bioscience Opt EIA, catalog #555165). They were assayed in 4 independent experiments with 2-3 technical replicate wells.

### Quantification of cytokine expression by qPCR

Stimulated J774 macrophages were washed with PBS and transferred into 1 mL Trizol (Invitrogen) and RNA was extracted following the manufacturer’s protocol. Isolated RNA was subjected to DnaseI treatment (Invitrogen) prior to cDNA synthesis. cDNA synthesis was completed using qScript cDNA SuperMix (Quantabio). qPCR amplification was performed in triplicate with SYBR green qPCR chemistry (Bioline) using primers for *Tnfa* (F-5’- ACCTCACACTCAGATCATCTTCTC-3’, R-5’- TGAGATCCATGCCGTTGG-3’), *Il1b* (F-5’- ACAGAATATCAACCAACAAGTGATATTCTC-3’, R-5’- GATTCTTTCCTTTGAGGCCCA-3’), *Ifnγ*(F-5’- CGGCACAGTCATTGAAAGCC-3’, R-5’- TGTCACCATCCTTTTGCCAGT-3’), *Csf2* (F-5’- ATGCCTGTCACGTTGAATGAAG-3’, R-5’- GCGGGTCTGCACACATGTTA-3’), *Il12b(p40)* (F-5’- CGCAAGAAAGAAAAGATGAAGGAG-3’, R-5’- TTGCATTGGACTTCGGTAGATG-3’), and *Gapdh* (F-5’- GGTGAAGGTCGGAGTCAACGGA-3’, R-5’- GAGGGATCTCGCTCCTGGAAGA-3’) on a QuantStudio 6 Real-Time PCR System. *C_t_* values were normalized to *Gapdh* to generate Δ*C_t_* values, and fold changes were calculated by ΔΔ*C_t_* to the Δ*C_t_* of unstimulated controls. These were assayed in 3 independent experiments in technical triplicates.

### Quantification of strains by qPCR

Stool DNA (6 ng each) from three mice was subjected to qPCR with primer pairs targeting each barcode to quantify relative amounts of each barcoded strain from within a complex population. Amplification was performed in duplicate using SYBR green qPCR chemistry (Bioline) using a universal barcode forward primer (F-5’- GCTTGGTTAGAATGGGTAACTAGTTTGCAG-3’) and barcode-specific reverse primers for A1 (R-5’- TTCCCGAGCGCACCACAAA-3’), A3 (R-5’- ACACATACATCTCGCACGCAAACG-3’), B6 (R-5’- AAACCAACATCTCCCTCGCCC-3’), C2 (R-5’- GGTGATGTTTGGGCGTGGTAGAA-3’), C5 (R-5’- ATAACACTCCCGCCCACGAGAA-3’), D2 (R-5’- TTCGAACTCGACCGCCAACCAAA-3’), and D5 (R-5’- CCACTCAATCACGCAACACCC-3’) with *E. coli* 16S primers (F-5’- ATTGACGTTACCCGCAGAAGA-3’, R-5’- GGGATTTCACATCCGACTTGA-3’) [34–35] on a QuantStudio 6 Real-Time PCR System. *C_t_* values were normalized to *E. coli* 16S rRNA to generate Δ*C_t_* values, and fold changes were calculated by ΔΔ*C_t_* to the Δ*C_t_* of the mouse inoculum.

### Mock tissue-associated microbiome

Aliquots of purified genomic DNA isolated from independent strain cultures were mixed at equal abundance by weight. HMW DNA was isolated from a frozen aliquot of 10^7^ HEK293 cells using the Circulomics Nanobind kit (https://www.circulomics.com/nanobind) per the manufacturer’s recommended protocol. Human DNA was mixed with microbial mixture at a ratio of 99.5: 0.5% to represent a realistic tissue-associated microbiota composition.

### Full-length rRNA amplicon sequencing

For stool and *in vitro* host/microbiota samples, primers for proximal 16S (27F: AGRGTTTGATYHTGGCTCAG) and distal 23S (2241R: ACCRCCCCAGTHAAACT) were used to amplify the full-length rRNA operon (4,500bp). Starting with ~5 ng expected microbial DNA, 25 ul PCR reactions were prepared with LongAmp Taq 2x Master Mix (New England Biolabs, Inc; M0287L) and 0.4 uM of each primer. We ran 20 cycles consisting of denaturation at 94°C for 10s, annealing at 51°C for 30s, and extension at 65°C for 225s, followed by final extension at 65°C for 10min. Typical yield is 700ng of full-length amplicons, which were verified by agarose gel electrophoresis and fluorescent quantification (Qubit). Amplicon libraries were prepared for nanopore sequencing using the Oxford Nanopore Ligation Sequencing Kit (LSK109) per the manufacturer’s protocol and sequenced on R9.4.1 flow cell on GridION with real-time basecalling using Guppy v3.2.2 in high-accuracy mode.

## Results

### Genome assembly and annotation

To establish a baseline for variation among known enteric *E. coli*, we produced finished genomes for eight AIEC and non-AIEC strains (NC101 [3,12], CU39ES-1-A1, CU532-9-A3, CU568-3-C5, CU37RT-2-D2, CU42ET-1-D5, HM670-C2, and LF82-B6). These genomes were sequenced, assembled, polished, and annotated as described in the Methods. Sequence data and assembled genomes have been made publicly available under NCBI BioProject ID PRJNA759208. Each genome was assembled into a single complete, circular contig representing the entire genome (Table 2). Table 3 describes the serotype (O and H), multi-locus sequence type (MLST), and virulence and antibiotic genes found in each. Figure 2 illustrates these eight genomes, annotated genes, and approximate homology between them.

**Fig 2.**
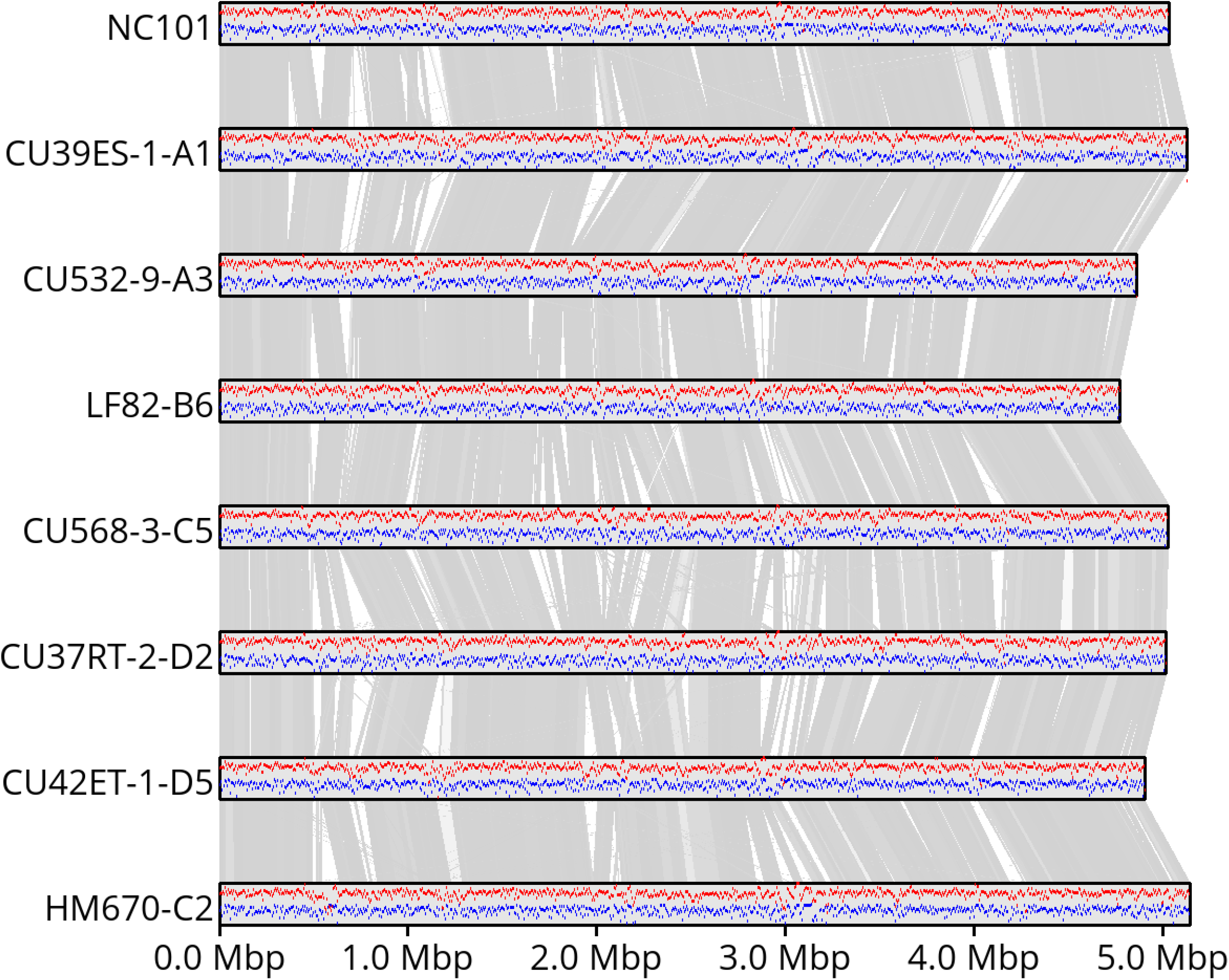
Assembled genomes, illustrating relative GC content (red) and gene content (blue) for each strain, and approximate synteny between assemblies represented in gray, where darker bars represent higher identity (darkest gray is ~100% identity, white is <70%).

**Table 2.** Genome size and number of genes identified in assembled E. coli genomes.

| Strain | Genome size | Genes |
| --- | --- | --- |
| HM670-C2 | 5,141,716 | 4,762 |
| CU39ES-1-A1 | 5,124,020 | 4,630 |
| CU532-9-A3 | 4,858,260 | 4,433 |
| LF82-B6 | 4,772,884 | 4,399 |
| CU568-3-C5 | 5,024,072 | 4,681 |
| CU37RT-2-D2 | 5,015,239 | 4,632 |
| CU42ET-1-D5 | 4,903,963 | 4,438 |
| NC101 | 5,030,087 | 4,630 |

**Table 3.** Characterization of each strain by serotype and MLST, including virulence, and antibiotic resistance genes.

| Strain | Serotype | MLST | Virulence genes | Antibiotic resistance |
| --- | --- | --- | --- | --- |
| HM670-C2 | O2:H1 | 73 | clbB*, pic*, iha, yfcV*, sat, ompT*, kpsMII, sitA, usp, fyuA*, papA_F43*, vat, tcpC, terC*, papA_F7-1, gad, iucC*, iutA, kpsE, chuA, iss, papC*, irp2, ireA | mdf(A)*, sitABCD* |
| CU39ES-1-A1 | O25:H1 | 73 | clbB, papA_F14, kpsMII_K5, mchB, mchC*, mchF, iha, yfcV*, sat, ompT*, mcmA, sitA, usp, fyuA*, iutA, hra, vat*, pic, cea, terC*, focI, iroN, gad, iucC, focG*, kpsE, focCsfaE, chuA*, iss, papA_F43*, hlyA, papC*, irp2 | mdf(A)*, sitABCD* |
| CU532-9-A3 | O4:H5 | 12 | ompT*, clbB, chuA, iss, sitA, usp*, terC*, tcpC, yfcV*, irp2, fyuA*, gad, vat*, kpsE* | mdf(A)*, sitABCD* |
| LF82-B6 | O83:H1 | 135 | ompT*, kpsE*, chuA, iss, sitA, usp*, terC*, ibeA, yfcV*, irp2*, fyuA*, kpsMII_K5, gad, vat | mdf(A)*, sitABCD* |
| CU568-3-C5 | O25:H4 | 131 | ompT*, kpsE, gad, chuA, iss, sitA*, usp*, terC*, iutA, papA_F43*, irp2, fyuA*, kpsMII_K5, iha, iucC, yfcV*, sat | mdf(A)*, sitABCD* |
| CU37RT-2-D2 | O16:H6 | 144 | ompT*, kpsE, gad*, chuA*, sitA*, usp*, terC*, papA_F7-1*, iutA, neuC, tcpC, kpsMII_K1, papC*, yfcV*, irp2*, fyuA*, iha*, iucC*, vat*, sat* | mdf(A)*, sitABCD* |
| CU42ET-1-D5 | O99:H6 | 1859 | papA_F12*, kpsMII_K1*, mchB, mchC*, mchF*, ibeA*, yfcV*, ompT*, sitA*, usp*, fyuA*, hra, vat*, pic*, terC*, focI*, sfaE*, iroN*, gad*, focG*, kpsE, chuA*, papC*, neuC, irp2* | mdf(A)*, sitABCD* |
| NC101 | O2:H6 | 998 | clbB, kpsMII_K1, pic*, ibeA, yfcV*, ompT*, sitA, usp*, fyuA*, vat, tcpC, sfaS, terC*, focI*, iroN*, gad*, kpsE, focCsfaE*, chuA, iss, neuC, irp2 | mdf(A)*, sitABCD* |

### Strain-level variation

As illustrated in Figure 2, we observe significant genome-wide structural variation while broadly conserving GC content and annotated gene density within homologous blocks, even among these closely-related strains with similar adherent-invasive (AI) or non-AI phenotype. We computed the pairwise hamming distance [37] between sequences across each strain and each copy of the rRNA operon for the 16S, ITS, 23S, and entire rRNA region (Figure 3).

**Fig 3.**
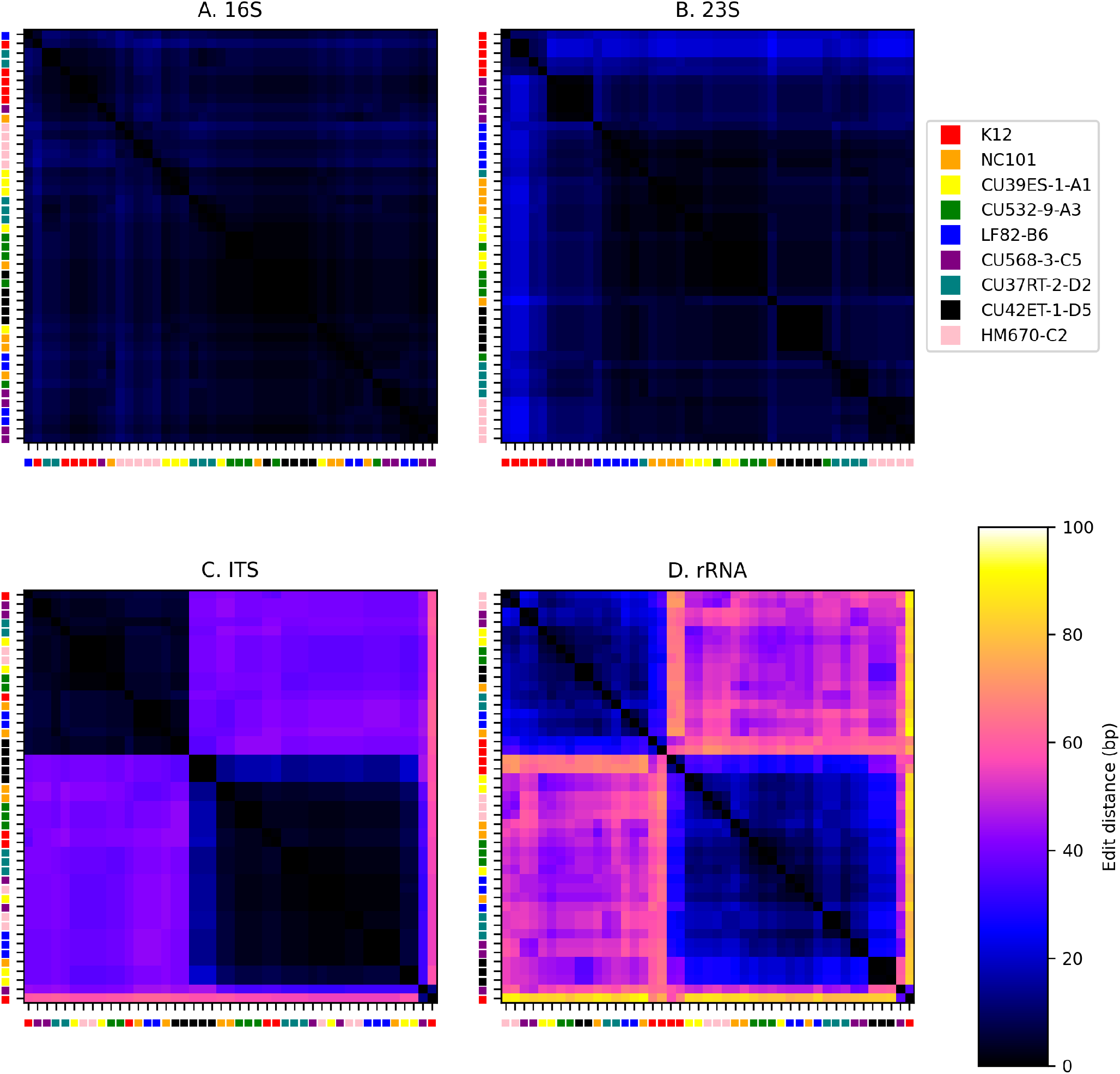
Difference between 16S (a), 23S (b), ITS (c), and entire rRNA (d) sequences across strains and copies. Heatmap shows the relative number of discriminating sites (alleles) between sequences. Sequences are clustered hierarchically using Ward’s variance minimization and are labeled by strain.

We evaluated several regions independently for informative variation among these strains *in silico* by aligning our whole-genome sequence to either 1) the V1-V2 hypervariable region of 16S, 2) V3-V4 hypervariable region, 3) the entire 16S sequence containing all nine hypervariable regions and conserved spacers, 4) the entire rRNA operon, including 16S, ITS, and 23S, and 5) the entire genome. Very long nanopore reads allow us to use this as a proxy for PCR amplification of the various rRNA amplicon analyses since these reads average 10-20 Kbp (see Table 4), and thereby most often cover the entire region of interest.

**Table 4.** Nanopore sequencing statistics for each strain, after demultiplexing.

| Strain | # reads | Total Gbp | Average read length | N50 |
| --- | --- | --- | --- | --- |
| HM670-C2 | 129,071 | 3.26 | 25,252 | 53,682 |
| CU39ES-1-A1 | 33,058 | 1.30 | 39,369 | 76,514 |
| CU532-9-A3 | 94,599 | 1.46 | 15,384 | 54,555 |
| LF82-B6 | 126,340 | 3.19 | 25,252 | 55,067 |
| CU568-3-C5 | 239,866 | 3.73 | 15,547 | 48,832 |
| CU37RT-2-D2 | 61,344 | 2.14 | 34,912 | 58,392 |
| CU42ET-1-D5 | 112,086 | 2.90 | 25,871 | 56,055 |
| NC101 | 22,886 | 0.44 | 19,341 | 78,877 |

In general, and not unexpectedly, the larger the region used for classification, the greater the accuracy in detecting the correct strain (Table 5). Hypervariable regions V1-V2 (27F-338R) and V3-V4 (343F-806R) produces poor results (averaging 35% and 27% accuracy, respectively), little better than chance since there is little or no discriminating variation within that region. The entire 16S gene performs better (70%). Notably, there is a huge amount of variation in the accuracy across strains owing to their relative dissimilarity in a particular region. Full-length rRNA achieves an average 87.3% accuracy at the strain level, followed closely by all reads aligned against the entire genome (i.e. shotgun metagenomics) at 91%. Additional benefits of whole-metagenome sequencing include the observation of genes that may imply the functional capacity of the community without explicitly characterizing its taxonomic structure, and the ability to assemble the metagenome into an approximation of its component microbial genomes. These are important, tangible benefits, but are beyond the scope of this paper.

**Table 5.** Classification accuracy using commonly amplified 16S hypervariable regions, V1-V2 and V3-V4, the entire 16S and rRNA operons, and shotgun sequencing of the entire genome. While short hypervariable regions perform poorly and inconsistently across these samples, classification based on the entire rRNA operon (~4,500bp) approaches classification accuracy of whole-genome sequencing.

| Strain | V1-V2 | V3-V4 | 16S | rRNA | Genomic |
| --- | --- | --- | --- | --- | --- |
| HM670-C2 | 14.48 | 29.66 | 61.95 | 71.75 | 74.85 |
| CU39ES-1-A1 | 14.72 | 12.50 | 47.39 | 70.73 | 84.79 |
| CU532-9-A3 | 17.52 | 19.61 | 96.93 | 93.44 | 90.44 |
| LF82-B6 | 93.18 | 18.66 | 97.62 | 87.07 | 94.78 |
| CU568-3-C5 | 89.62 | 87.39 | 99.06 | 99.19 | 97.09 |
| CU37RT-2-D2 | 14.68 | 31.25 | 59.16 | 96.83 | 97.57 |
| CU42ET-1-D5 | 17.60 | 2.90 | 55.26 | 99.07 | 96.13 |
| NC101 | 16.67 | 10.77 | 45.71 | 80.18 | 93.43 |
| *Average* | *34.81* | *26.59* | *70.39* | *87.28* | *91.14* |

There is a relatively small improvement in classification accuracy using the entire genomes compared to the full-length rRNA operon. Like commonly used 16S primers that target subsets of the hypervariable regions, the primer pair used for the full operon are well-conserved across bacteria, but presents a much larger sequence with which to accurately classify sequences. This coupled with cost-effective multiplexing and long-read sequencing, despite much higher error rates than standard Illumina sequencing, should make this a viable approach for characterizing complex microbiome samples.

### In vivo colonization studies

We performed *in vivo* colonization studies using a well-established IBD mouse model [3, 12–13, 36, 38–39]. We first determined if these human-derived strains and murine-derived NC101 could colonize and persist in mice in the presence of a competing microbiota. We colonized two singly-housed, adult, germ-free, *Il10*^-/-^ mice each with a single *E. coli* strain, then after one week provided niche competition by gavaging with a murine fecal transplant as outlined in Figure S2. After five weeks total, gastrointestinal tissues, including ileal tissue, cecal content, colon content, colon tissue, and colon mucus layer, were harvested and viable *E. coli* were quantified by serial plating. Table S1 reveals that almost all *E. coli* strains could colonize the lumen and intestinal tissue with complex community competition, even without kanamycin to suppress the competing microbiota. Two strains of *E. coli* colonized inconsistently across both mice, with high levels of *E. coli* in one mouse and low or undetected levels of *E. coli* in the other. To verify if this would persist across a larger cohort, consistent colonizing strain, CU37RT-2-D5, and inconsistent colonizing strain, HM670-C2, were given by gavage to seven or eight germ-free, *Il10*^-/-^ mice, respectively. After one week we provided niche competition by gavaging with a murine fecal transplant, and then after three weeks we provided kanamycin water *ad libitum* for two weeks to supress the competing microbiota as outlined in Figure S3. After ten weeks total, gastrointestinal tissues, including colon content, colon tissue, and colon mucus layer, were harvested and viable *E. coli* were quantified by serial plating. Table S2 reveals that consistent colonizing strain CU37RT-2-D5 colonized all seven mice to high levels, while inconsistent colonizing strain HM670-C2 colonized five of eight mice at the lumen and intestinal tissue. Future studies are needed to validate these results and draw strong conclusions, including colonization in inflamed and non-inflamed environments.

To validate the pro-inflammatory potential of these strains, each strain was co-cultured with J774 macrophages and inflammatory colitis-inducing cytokine transcription and secretion was measured [40–43]. The results in Figure 4 demonstrate that all strains induced inflammatory cytokine production. Inducing inflammatory cytokine production is an indicator of inflammatory potential, but it is not a known predictor of tissue inflammation or colonization *in vivo*. Further effort is needed to compare *in vitro* cytokine production with *in vivo* inflammation elicited by colonizing AIEC isolates.

**Figure 4:**
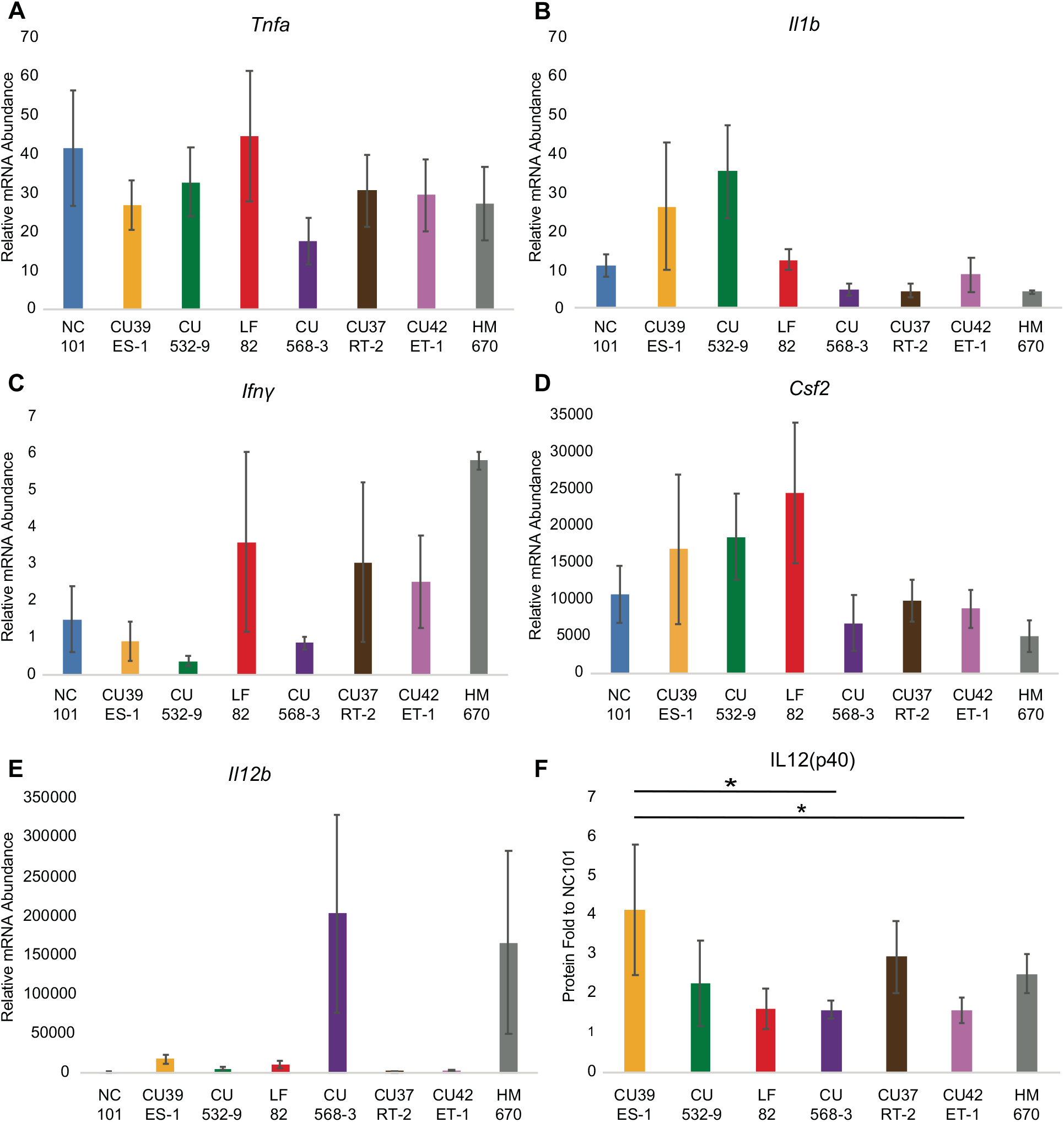
Inflammatory cytokine production by macrophages stimulated with clinical *E. coli* strains. (a-e) Relative mRNA abundance of each cytokine from stimulated J774 macrophages determined by qRT-PCR. Values normalized to gapdh, and fold increase relative to unstimulated controls. Each bar represents the mean across 3 independent experiments in triplicate and error bars show standard deviation (*p<0.05, Tukey’s multiple comparisons test). If not otherwise noted, there is no significance. f) Protein abundance of IL12(p40) from stimulated J774 macrophages determined by ELISA. Values normalized to NC101 stimulated macrophages. Each bar represents the mean across 4 independent experiments in duplicate and error bars show standard deviation (*p<0.05, Tukey’s multiple comparisons test). If not otherwise noted, there is no significance

### Full-length rRNA metataxonomics

To further evaluate the utility of performing strain-level metataxonomics using full-length rRNA sequencing on an Oxford Nanopore device, we prepared and sequenced one stool sample from *in vivo Il10*^-/-^ colonization studies depicted in Figure 5. *Il10*^-/-^ mice, a well-established model of microbially driven intestinal inflammation, were colonized with an equally-proportioned pool of the seven *E. coli* clinical isolates, and then housed in an SPF facility where they become colonized over time with a natural mouse microbiome [3, 36]. After ten weeks, the mice were sacrificed and stool samples were collected for nanopore sequencing and quantification of each strain by targeted qPCR. We extracted DNA from the resulting stool samples. Full-length rRNA amplification and sequencing using universal 16S 27F and 23S 2241R primers (see Methods) produced >600,000 reads with a median length of 4,149 bp (Table 6, Figure 6). We assigned these reads to individual rRNA copies by aligning to our database of known sequences in the assembled genomes (40 copies, five in each of eight strains). Figure 7 illustrates the distribution of alignment identity (number of matching nucleotides) over the full-length rRNA (4,245bp). Sequences derived from the stool sample show a bimodal distribution with peaks at 3.9Kbp (92% identity) and 2.9Kbp (68% identity), representing the expected divergence and sequencing error matching *E. coli* and non-*Escherichia* genera, respectively. The proportion of reads assigned to each strain is shown in Figure 5(b). There is a skewed representation of strains in the stool sample due to natural variation in their ability to colonize and thrive and due to interstrain competition between isolates in the *Il10*^-/-^ mouse gut. We confirmed the relative abundance of each strain in the stool sample through qPCR of the molecular barcode inserted in the neutral Tn7 site of each *E. coli* genome [34]. The qPCR showed a similar result, as there was skewing among the *E. coli* isolates in the mouse gut as shown in Figure 5 with the CU568-3-C5 strain most abundant. Both consistent and inconsistent colonizing strains showed lower abundance in this stool sample as there were multiple *E. coli* isolates providing competition upon initial infection, and this data is a measure of relative abundance across *E. coli* strains and not total load with the stool. Additionally, the inconsistent colonizing strain still colonized in 5 of 8 mice (Table S2).

**Figure 5.**
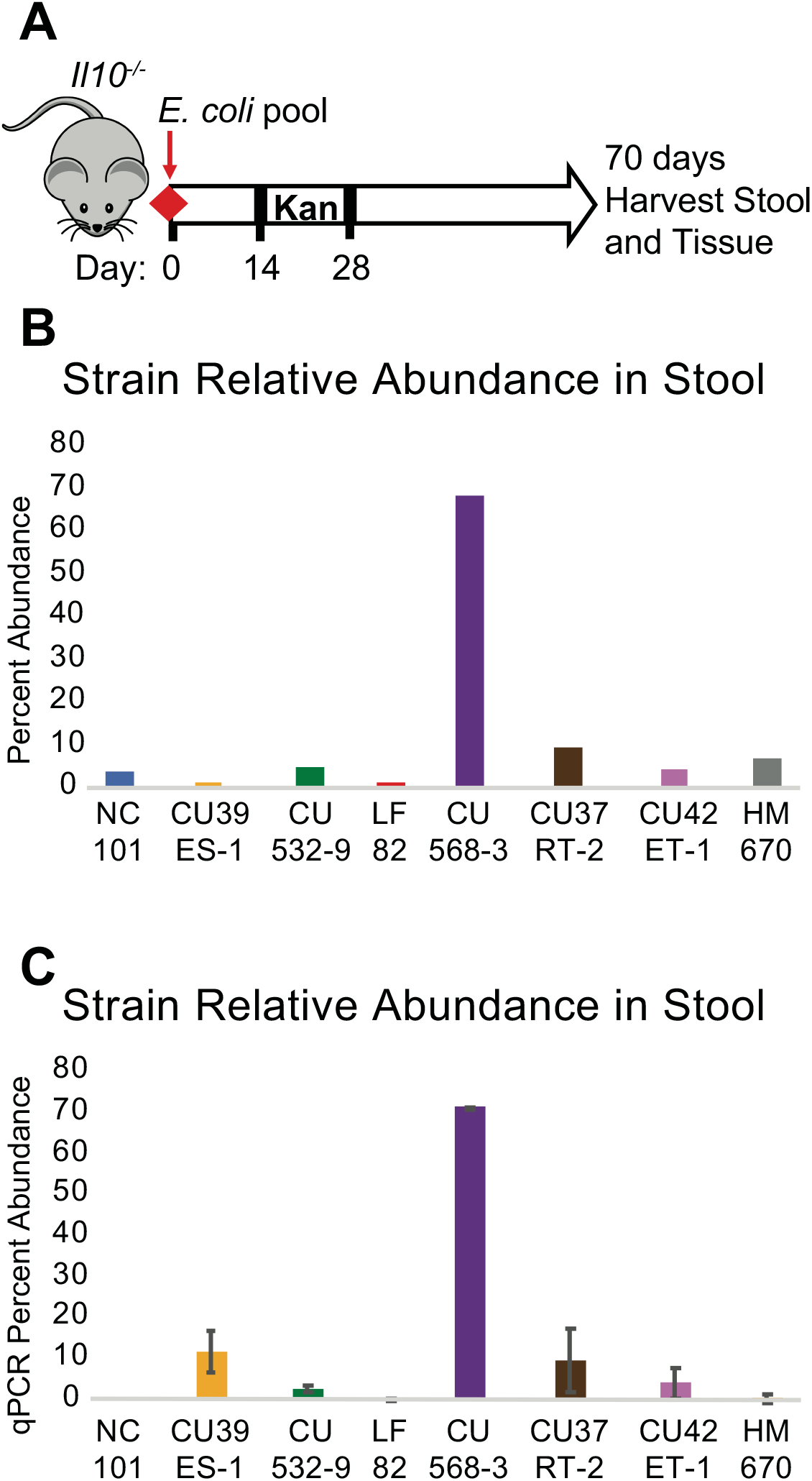
Strain distribution of seven E. coli isolates in murine stool sample. (a) Timeline for mouse experiment. (b) Relative percent abundance of each strain determined by full-length rRNA sequencing. Off-target hits to NC101 (4%) are expected to happen by chance given the classification error rate for the whole rRNA sequence (~13%, see Table 5). (c) Relative percent abundance of each strain determined by quantitative PCR of molecular barcode. Values normalized to *E. coli* 16S rRNA, and fold increase relative to pooled *E. coli* inoculum. Each bar represents the mean across 3 murine stool samples in duplicate and error bars show standard deviation.

**Figure 6.**
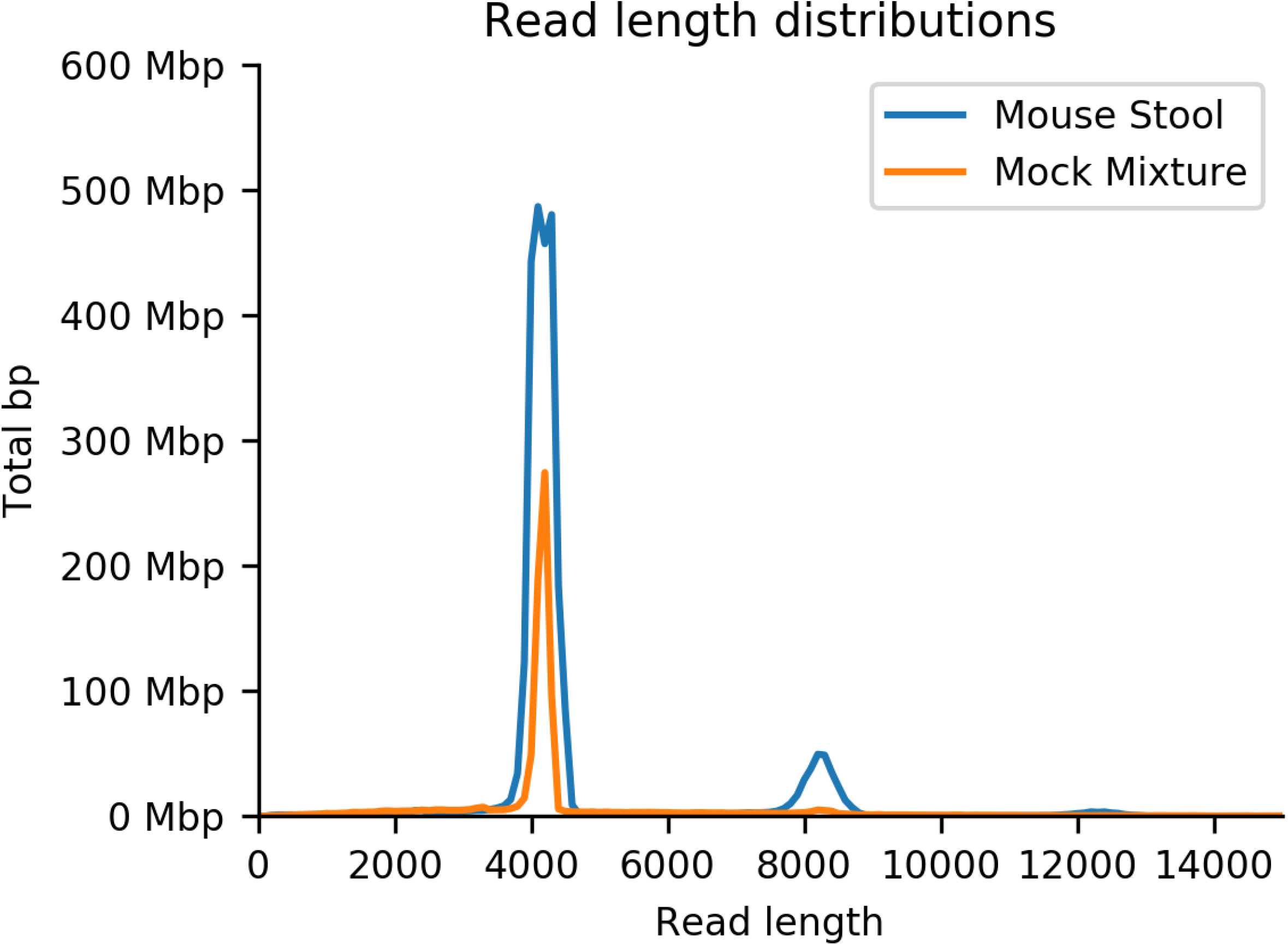
Read length distribution for stool and mock mixture full-length rRNA amplicon sequencing. The secondary peak at ~8Kbp represents chimeric reads resulting from nanopore sequencing that are separated prior to alignment.

**Figure 7.**
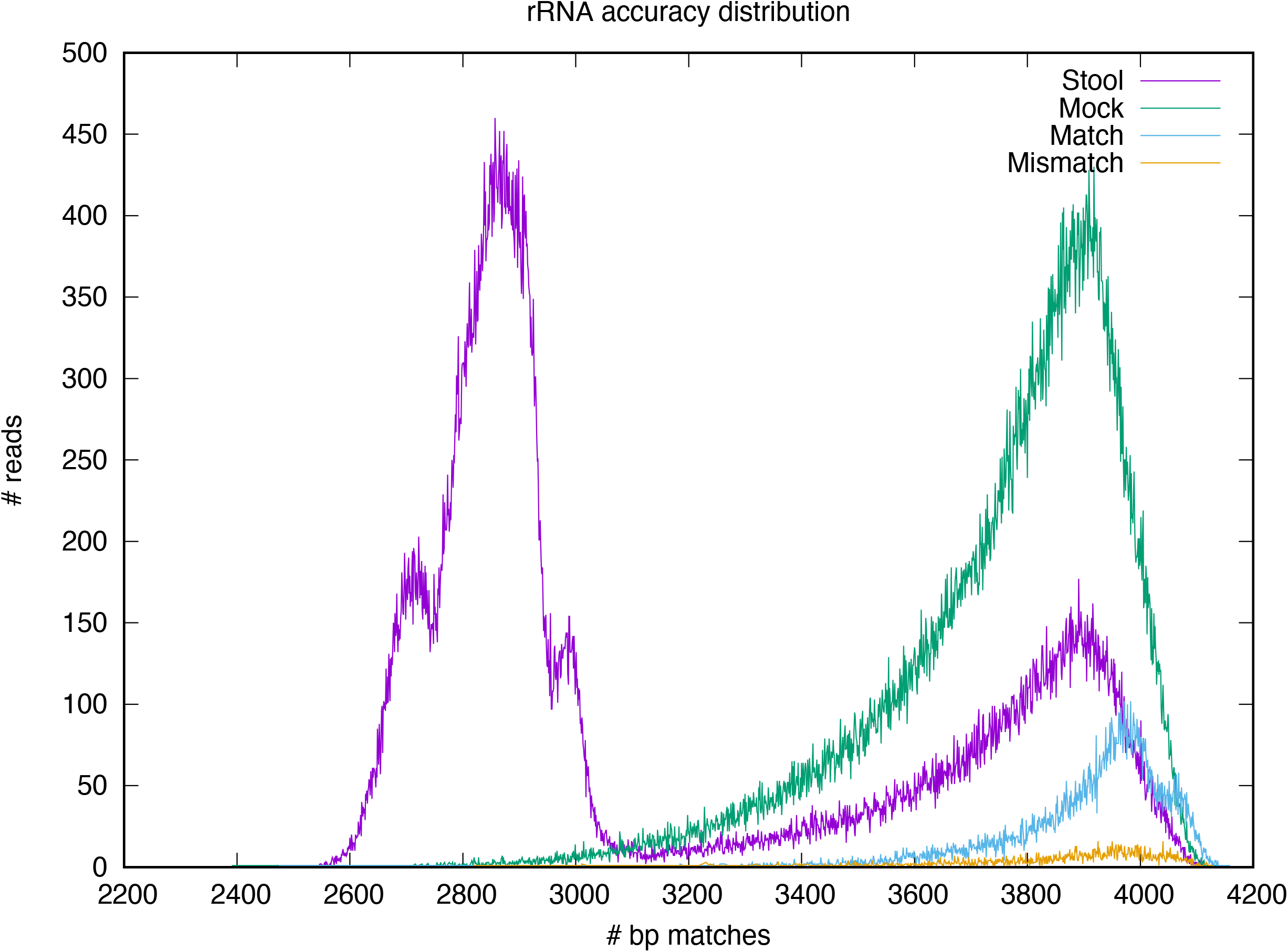
Distribution of nucleotide matches of the best alignment from amplified rRNA sequencing. Match/Mismatch represent the aligned rRNA segments from the barcoded whole-genome sequencing where we know the ground truth. Mismatches have a nearly identical distribution to matches, indicating sequencing errors caused miscategorization as a very closely-related rRNA. Mock shows a very similar distribution, but sequences from stool are bimodal indicating a (smaller) proportion of reads originated from these or similar E.coli rRNA, and a majority (~75%) originating from highly divergent sequences (70-80% identity) likely originating from non-Escherichia genera and phyla.

**Table 6.** Sequencing statistics for full-length rRNA amplicon sequencing.

| Sample | # reads | Total bp | Average read length | Median read length | Read N50 |
| --- | --- | --- | --- | --- | --- |
| Mouse stool sample | 657,234 | 2.78 Gbp | 4,229.65 | 4,149 | 4,194 |
| 0.5% *E.coli* mix, 99.5% HEK293 | 240,691 | 0.92 Gbp | 3,829.50 | 4,136 | 4,173 |

We additionally prepared a mock mixture consisting of all eight strains at even abundance mixed 0.5: 99.5% with human DNA (HEK293) to simulate a realistic tissue-associated microbiota sample. Full-length rRNA sequences were successfully amplified and sequenced as done with the stool sample. We observed relatively uniform representation of the eight strains in the mock mixture, demonstrating our ability to amplify and properly classify closely-related rRNA sequences in a host-tissue-like context, as illustrated in Figure 8.

**Figure 8.**
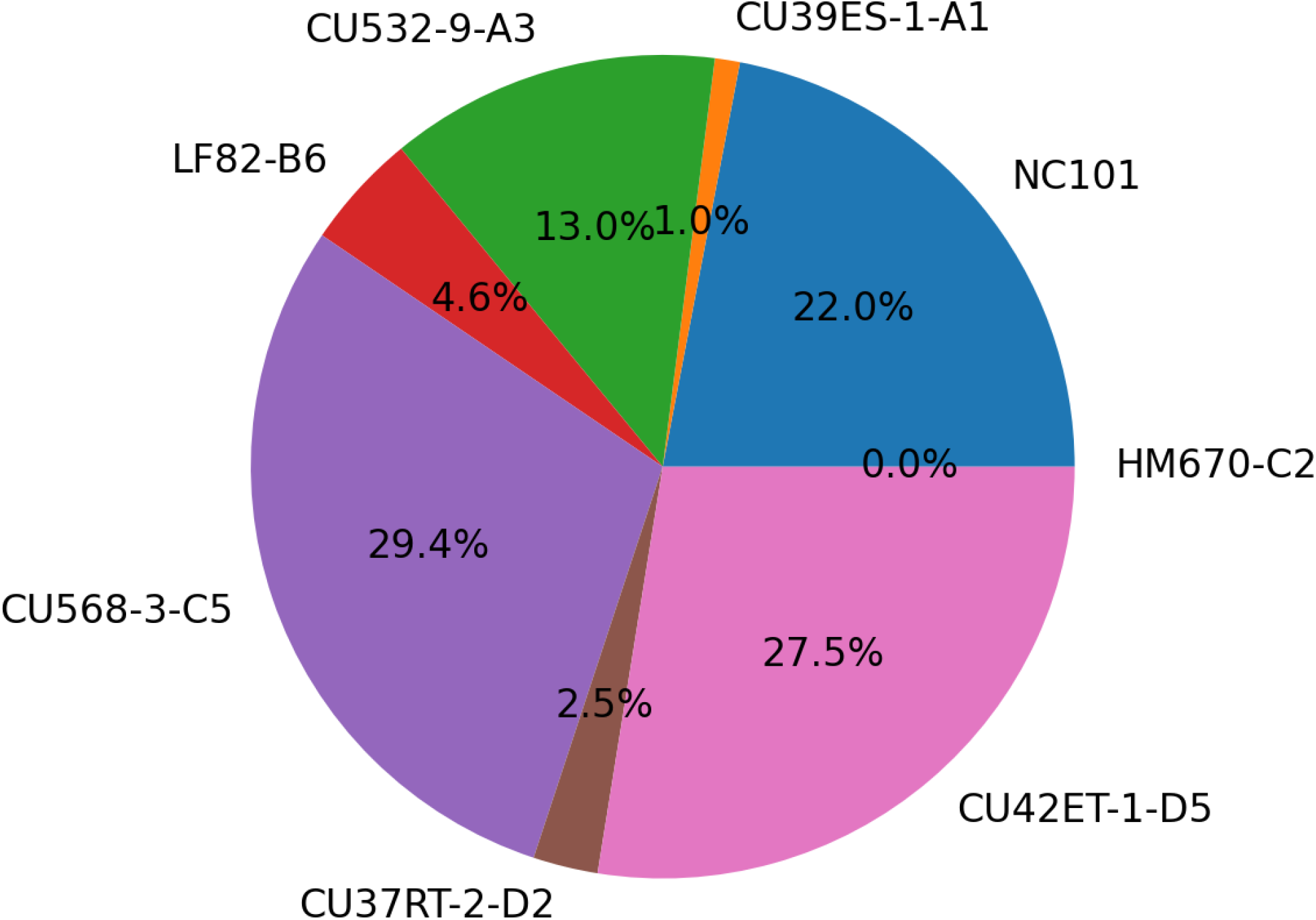
Strain representation based on full-length rRNA sequencing from mock host:microbial mixture. Mock mixture contained an approximately uniform quantify of all eight strains before amplification.

## Discussion

Alterations in the composition and function of gut microbial communities are implicated in both IBD and CRC, with CD patient microbiomes harboring an increase in mucosa-associated Enterobacteriaceae, including AIEC [7]. After excluding genes associated with diarrheagenic *E. coli*, AIEC are defined by their *in vitro* adherent-invasive behavior. There is a clear need to extend our knowledge of AIEC to pathoadaptive determinants of colonization and virulence *in vivo*. Recent work highlights mechanisms AIEC have evolved to more effectively colonize the host mucosa including hypermotility and metabolic changes [44] and T4SS-linked biofilm formation [45]. These efforts involved either long *in vivo* host adaptation experiments or genome-wide screens with a limited number of *E. coli* strains. Other molecular mechanisms including the type 1 pili-CEACAM6 interaction that promotes adhesion of AIEC to intestinal epithelial cells [46–47] do not offer a genetic marker to distinguish AIEC. Competitive colonization with pools of known AIEC and non-AIEC strains as a parallel strategy could help elucidate traits and genes important for colonization, but requires methods to discriminate between closely-related microbes. To relate specific traits of different AIEC to pre-clinical models and patients, we need to be able to distinguish individual strains from within a complex community. As AIEC are mucosa-associated, the ability to evaluate colonization of inflamed or cancerous lesions and normal mucosa would be optimal.

The commonly used metataxonomic approach of sequencing one or two amplified hypervariable regions of the 16S gene is unable to distinguish among closely-related, but functionally diverse, species and strains [21]. Consistent with previous work [17–21], we demonstrate that sequencing of longer regions of the 16S, ITS, and 23S operons, enabled by modern long-read sequencing technologies, dramatically increases taxonomic specificity. We illustrated the increase in sensitivity and specificity of strain detection using full-length rRNA amplification and whole-metagenome shotgun sequencing in *in silico* models, mock mixtures, and *in vivo* colonized murine microbiome.

This focused study highlights specifically the challenges in resolving closely-related but functionally divergent AIEC and non-adherent/invasive *E. coli* strains within mixed microbial populations. We demonstrated accurate identification and discrimination of these strains in complex *in vitro* and *in vivo* models relevant to colitis. Our novel nanopore sequencing approach illustrates an efficient and effective method for discrimination of known pathogenic or specifically colonized microbiota, including tissue-associated bacteria, to enable investigation of spatiotemporal dynamics of microbial colonization without relying on genetic engineering, molecular barcoding [34], or *in vivo* fluorescence markers [48].

While these results suggest a method for sensitive identification of known strains, this approach does not suggest a general taxonomic analysis protocol for microbiome studies since the genomes of the strains of interest must be known *a priori* in order to properly identify them. Full-length rRNA sequencing following PCR amplification permits accurate strain identification even when the microbial abundance - for example in tissue-associated microbial samples - is very low. We demonstrated high concordance among relative abundance between sequenced full-length rRNA sequences and qPCR results targeting the inserted molecular barcodes, suggesting that abundances are reliably preserved through amplification and nanopore sequencing. However, even higher accuracy and explicit identification of microbial genic content can be captured by shotgun metagenome sequencing when there exists enough material to extract high-quality microbial DNA.

Future work includes optimization of the full-length rRNA protocol for tissue samples with adherent microbes. Our analysis of *in vitro* mixtures indicate this should be feasible with microbial:host DNA content as low as 0.5% - consistent with expected abundances [49–50] in many human mucosa, including the gut - but we have so far failed to amplify full-length rRNA from resected mouse gut. Additionally, given the extraordinary utility of shotgun metagenomic sequence, it is worthwhile to explore the several existing methods for selective extraction and sequencing of microbial DNA to expand the availability of non-PCR-based approaches in the context of tissue-associated microbial communities.

## Acknowledgements

This work was supported by the following grants: NIH/NIDDK K01 DK119582 (JW), SPIRE (NIH IRACDA program) NIH/NIGMS K12 GM000678-21 (RMB), NIH/NIDDK K01 DK103952 (JCA), NIH/NIDDK R01 DK124617 (JCA), NIH/NIAID R21 AI159786 (JCA), American Gastroenterological Association Augustyn Award in Digestive Cancer (JCA), Lineberger Comprehensive Cancer Center Pilot Grant (JCA).

We acknowledge the Gnotobiotic Core at the UNC Center for Gastrointestinal Biology and Disease (CGIBD: supported by NIH P30DK34987), the National Gnotobiotic Rodent Resource Center (supported by NIH P40 OD01995), and UNC’s High-Throughput Sequencing Facility. We also acknowledge Lacey Lopez, Christopher Broberg, Adrienne Franks, and Cassandra Barlogio for experimental assistance.

---

**S1 Figure.**
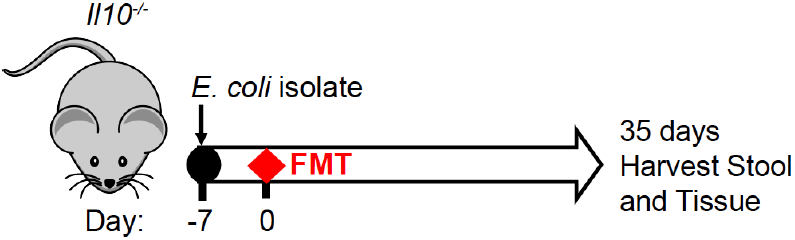
Timeline for mouse experiment to determine colonize of individual *E. coli* isolates.

**S1 Table.** *E. coli* isolates colonize the lower gastrointestinal tract of *Il10*^-/-^ mice with FMT competition. Values indicate CFUs/10 mg of tissue.

| Strain | Mouse Gender | Ileal Tissue | Cecal Content | Colon Content | Mucus Layer | Colon Tissue |
| --- | --- | --- | --- | --- | --- | --- |
| CU39ES-1/A1 | M | 65957 | 3916667 | 8588235 | 3556 | 12889 |
| CU39ES-1/A1 | F | 611 | 5882353 | 34166667 | 6500 | 10833 |
| CU532-9/A3 | M | 0 | 5806452 | 7058824 | 6667 | 455556 |
| LF82/B6 | M | 1875 | 232609 | 257732 | 200 | 1800 |
| LF82/B6 | F | 500 | 111111 | 74684 | 417 | 417 |
| CU568-3/C5 | M | 1250 | 40909091 | 5306122 | 24545 | 681818 |
| CU568-3/C5 | F | 0 | 0 | 71 | 0 | 0 |
| CU37RT-2/D2 | M | 0 | 0 | 0 | 0 | 0 |
| CU42ET-1/D5 | F | 28947 | 167857 | 320513 | 200 | 5133 |
| CU42ET-1/D5 | M | 107857 | 1333333 | ND | 1500 | 32308 |
| HM670/C2 | F | 0 | 34 | 0 | 0 | 188 |
| HM670/C2 | M | 13810 | 460317 | 422018 | 6583 | 25000 |
| NC101 | M | 500 | 39333 | 24615 | 0 | 70000 |

\*Mice were all singly-housed.
\* CU532-9/A3 F (dead), CU37RT-2/D2 F (dead), NC101 F (dead)

**S2 Figure.**
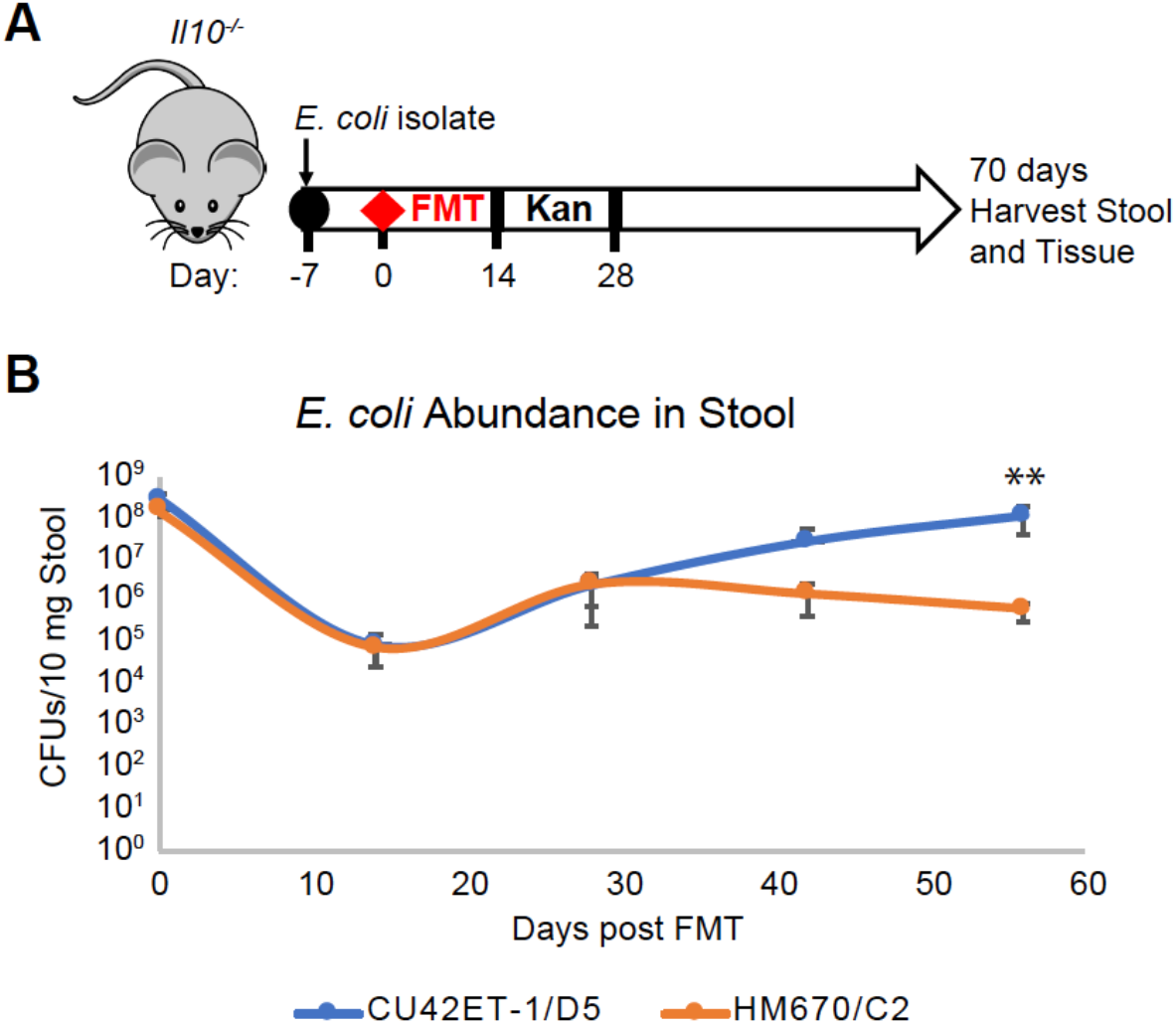
A. Timeline for mouse experiment for colonization of CU42ET-1/D5 or HM670/C2. B. Levels of *E. coli* in stool over time (n=7 in CU42ET-1/D5 and n=8 in HM670/C2). Data point indicates the mean and error bars indicates the standard deviation (**p<0.01 Mann-Whitney test). If not otherwise noted, there is no significance.

**S2 Table.** Consistent *E. coli* colonizer (CU42ET-1/D5) and inconsistent *E. coli* colonizer (HM670/C2) colonize the lower gastrointestinal tract of *Il10*^-/-^ mice with FMT competition.

| Mouse | Cage | Colon Content | Mucus Layer | Colon Tissue |
| --- | --- | --- | --- | --- |
| CU42ET-1/D5 1 | 1 | 19,083,969 | 400,000 | 1,181,818 |
| CU42ET-1/D5 2 | 1 | 7,565,217 | 177,778 | 722,222 |
| CU42ET-1/D5 3 | 1 | 3,802,817 | 5,455 | 150,000 |
| CU42ET-1/D5 4 | 2 | 49,494,949 | 210,526 | 894,737 |
| CU42ET-1/D5 5 | 2 | 37,419,355 | 67,308 | 730,769 |
| CU42ET-1/D5 6 | 2 | 33,750,000 | 225,000 | 616,667 |
| CU42ET-1/D5 7 | 2 | 95,890,411 | 45,000 | 4,000,000 |
| HM670/C2 1 | 3 | 553,191 | 333 | 66,667 |
| HM670/C2 2 | 3 | 1,193,277 | 7,250 | 300,000 |
| HM670/C2 3 | 3 | 2,019,231 | 0 | 17,000 |
| HM670/C2 4 | 3 | 2,179,487 | 1,125 | 13,125 |
| HM670/C2 5 | 4 | 204,301 | 1,000 | 10,778 |
| HM670/C2 6 | 4 | 0 | 0 | 0 |
| HM670/C2 7 | 4 | 139 | 0 | 0 |
| HM670/C2 8 | 4 | 0 | 0 | 0 |

## References

1. Sartor RB, Wu GD. Roles for Intestinal Bacteria, Viruses, and Fungi in Pathogenesis of Inflammatory Bowel Diseases and Therapeutic Approaches. Gastroenterology. 2017;152(2):327–339.e4.

2. Kappelman MD, Farkas DK, Long MD, Erichsen R, Sandler RS, Sørensen HT, et al. Risk of cancer in patients with inflammatory bowel diseases: A nationwide population-based cohort study with 30 years of follow-up evaluation. Clin Gastroenterol Hepatol [Internet]. 2014;12(2):265–273.e1. Available from: http://dx.doi.org/10.1016/j.cgh.2013.03.034

3. Arthur JC, Perez-Chanona E, Mühlbauer M, Tomkovich S, Uronis JM, Fan T-J, et al. Intestinal Inflammation Targets Cancer-Inducing Activity of the Microbiota. Science. 2012;338(6103):120–123.

4. Dejea CM, Fathi P, Craig JM, Boleij A, Taddese R, Geis AL, et al. Patients with familial adenomatous polyposis harbor colonic biofilms containing tumorigenic bacteria. Science. 2018;359(6375):592–7.

5. Darfeuille-Michaud A, Boudeau J, Bulois P, Neut C, Glasser A-L, Barnich N et al. High Prevalence of Adherent-Invasive Escherichia Coli Associated With Ileal Mucosa in Crohn’s Disease. Gastroenterology. 2004;127(2):412–21.

6. Martin HM, Campbell BJ, Hart CA, Mpofu C, Nayar M, Singh R, et al. Enhanced Escherichia coli adherence and invasion in Crohn’s disease and colon cancer. Gastroenterology. 2004;127(1):80–93.

7. Baumgart M, Dogan B, Rishniw M, Weitzman G, Bosworth B, Yantiss R, et al. Culture independent analysis of ileal mucosa reveals a selective increase in invasive Escherichia coli of novel phylogeny relative to depletion of Clostridiales in Crohn’s disease involving the ileum. ISME J. 2007;1(5):403–18.

8. Knights D, Lassen KG, Xavier RJ. Advances in inflammatory bowel disease pathogenesis: Linking host genetics and the microbiome. Gut. 2013;62(10):1505–10.

9. Frank DN, St. Amand AL, Feldman RA, Boedeker EC, Harpaz N, Pace NR. Molecular-phylogenetic characterization of microbial community imbalances in human inflammatory bowel diseases. Proc Natl Acad Sci U S A. 2007;104(34):13780–5.

10. Rooks MG, Veiga P, Wardwell-Scott LH, Tickle T, Segata N, Michaud M, et al. Gut microbiome composition and function in experimental colitis during active disease and treatment-induced remission. ISME J. 2014;8:1403–1417.

11. Vila AV, Imhann F, Collij V, Jankipersadsing SA, Gurry T, Mujagic A, et al. Gut microbiota composition and functional changes in inflammatory bowel disease and irritable bowel syndrome. Sci Trans Med. 2018;10(472):eaap8914.

12. Kim SC, Tonkonogy SL, Albright CA, Tsang J, Balish EJ, Braun J, et al. Variable phenotypes of enterocolitis in interleukin 10-deficient mice monoassociated with two different commensal bacteria. Gastroenterology. 2005;128(4):891–906.

13. Kim SC, Tonkonogy SL, Karrasch T, Jobin C, Balfour Sartor R. Dual-association of gnotobiotic IL-10-/-mice with 2 nonpathogenic commensal bacteria induces aggressive pancolitis. Inflamm Bowel Dis. 2007;13(12):1457–66.

14. Carvalho FA, Koren O, Goodrich JK, Johansson MEV, Nalbantoglu I, Aitken JD, et al. Transient inability to manage proteobacteria promotes chronic gut inflammation in TLR5-deficient mice. Cell Host Microbe. 2012;12(2):139–52.

15. O’Brien CL, Bringer MA, Holt KE, Gordon DM, Dubois AL, Barnich N, et al. Comparative genomics of Crohn’s disease-Associated adherent-invasive Escherichia coli. Gut. 2017;66(8):1382–9.

16. Dogan B, Scherl E, Bosworth B, Yantiss R, Altier C, McDonough PL, et al. Multidrug Resistance Is Common in Escherichia coli Associated with Ileal Crohn’s Disease. Inflamm Bowel Dis. 2013;19(1):141–150.

17. Benítez-Páez A, Portune KJ, Sanz Y. Species-level resolution of 16S rRNA gene amplicons sequenced through the MinION™ portable nanopore sequencer. GigaScience. 2016;5(1). doi:10.1186/s13742-016-0111-z.

18. Cuscó A, Catozzi C, Viñes J, Sanchez A, Francino O. Microbiota profiling with long amplicons using Nanopore sequencing: full-length 16S rRNA gene and the 16S-ITS-23S of the rrn operon [version 2; peer review: 2 approved, 3 approved with reservations]. F1000Research 2019, 7:1755.

19. Callahan BJ, Wong J, Heiner C, Oh S, Theriot CM, Gulati AS et al. High-throughput amplicon sequencing of the full-length 16S rRNA gene with single-nucleotide resolution. Nucleic Acids Research. 2019;47(18):e103. doi:10.1093/nar/gkz569

20. Karst SM, Ziels RM, Kirkegaard RH, Sørensen EA, McDonald D, Zhu Q, et al. High-accuracy long-read amplicon sequences using unique molecular identifiers with Nanopore or PacBio sequencing. Nat Methods. 2021; 18:165–169. https://doi.org/10.1038/s41592-020-01041-y

21. Johnson JS, Spakowicz DJ, Hong B-Y, Petersen LM, Demkowicz P, Chen L, et al. Evaluation of 16S rRNA gene sequencing for species and strain-level microbiome analysis. Nat Commun. 2019;10(5029). doi:10.1038/s41467-019-13036-1.

22. Sambrook J, Russell DW. Purification of Nucleic Acids by Extraction With Phenol:chloroform. CSH Protoc, 2006 (1).

23. Tyson J. Rocky Mountain adventures in Genomic DNA sample preparation, ligation protocol optimisation / simplification and Ultra long read generation. protocols.io. 2020. dx.doi.org/10.17504/protocols.io.7euhjew.

24. Li H. Minimap and miniasm: fast mapping and de novo assembly for noisy long sequences. Bioinformatics. 2016;32(14):2103–2110.

25. Vaser R, Sović I, Nagarajan N, Šikić M. Fast and accurate de novo genome assembly from long uncorrected reads. Genome Res. 2017;27:737–46. doi:10.1101/gr.214270.116.

26. Oxford Nanopore Technologies. Medaka; 2018 [cited 2020 March 3]. Available from: https://nanoporetech.github.io/medaka/

27. Walker BJ, Abeel T, Shea T, Priest M, Abouelliel A, et al. Pilon: An Integrated Tool for Comprehensive Microbial Variant Detection and Genome Assembly Improvement. PLOS ONE. 2014;9(11):e112963. doi:10.1371/journal.pone.0112963.

28. Tatusova T, DiCuccio M, Badretdin A, Chetvernin V, Nawrocki EP, Zaslavsky L et al. NCBI prokaryotic genome annotation pipeline. Nucleic Acids Res. 2016;44(14):6614–24. doi: 10.1093/nar/gkw569.

29. Buchfink B, Xie C. Huson DH. Fast and sensitive protein alignment using DIAMOND. Nature Methods. 2015;12:59–60.

30. Inouye M, Dashnow H, Raven L-A, Schultz MB, Pope BJ, Tomita T et al. SRST2: Rapid genomic surveillance for public health and hospital microbiology labs. Genome Med. 2014;6(11):90.

31. Center for Genomic Epidemiology; 2011 [cited 2020 March 3]. [Internet]. Available from: http://www.genomicepidemiology.org/

32. Li H. Minimap2: pairwise alignment for nucleotide sequences. Bioinformatics. 2018;34(18):3094–3100.

33. Tegha G, Ciccone EJ, Krysiak R, Kaphatika J, Chikaonda T, Ndhlovu I, et al. Genomic epidemiology of Escherichia coli isolates from a tertiary referral center in Lilongwe, Malawi. Microbial Genomics. 2021;7(1):mgen000490. doi:10.1099/mgen.0.000490.

34. Walters MS, Lane MC, Vigil PD, Smith SN, Walk SR, Mobley HLT. Kinetics of Uropathogenic Escherichia Coli Metapopulation Movement During Urinary Tract Infection. mBio. 2012;3(1):e00303–11.

35. Gonzalez RJ, Lane MC, Wagner NJ, Weening EH, Miller VL. Dissemination of a highly virulent pathogen: tracking the early events that define infection. PLoS Pathog. 2015;11(1):e1004587.

36. Arthur JC, Gharaibeh RZ, Mühlbauer M, Perez-Chanona E, Uronis JM, McCafferty J et al. Microbial genomic analysis reveals the essential role of inflammation in bacteria-induced colorectal cancer. Nature Communications. 2014;5(4724). doi:10.1038/ncomms5724

37. Hamming RW. Error detecting and error correcting codes. The Bell System Technical Journal. 1950;29(2):147–160. doi: 10.1002/j.1538-7305.1950.tb00463.x.

38. Kühn R, Löhler J, Rennick D, Rajewsky K, Müller W. Interleukin-10-deficient mice develop chronic enterocolitis. Cell. 1993;75(2):263–74.

39. Ellermann M, Gharaibeh RZ, Fulbright L, Dogan B, Moore LN, Broberg CA et al. Yersiniabactin-Producing Adherent/Invasive *Escherichia coli* Promotes Inflammation-Associated Fibrosis in Gnotobiotic *Il10^-/-^* Mice. Infection and Immunity. 2019;87(11):1–18.

40. Sartor RB. Cytokines in intestinal inflammation: Pathophysiological and clinical considerations. Gastroenterology. 1994;106(2):533–9.

41. Patwa LG, Fan TJ, Tchaptchet S, Liu Y, Lussier YA, Sartor RB, et al. Chronic intestinal inflammation induces stress-response genes in commensal Escherichia coli. Gastroenterology. 2011;141(5):1842–1851.e10. doi:10.1053/j.gastro.2011.06.064

42. Ellermann M, Huh EY, Liu B, Carroll IM, Tamayo R, Sartor RB. Adherent-invasive Escherichia coli production of cellulose influences iron-induced bacterial aggregation, phagocytosis, and induction of colitis. Infect Immun. 2015;83(10):4068–80.

43. Segain JP, Galmiche JP, Raingeard De La Blétière D, Bourreille A, Leray V, Gervois N, et al. Butyrate inhibits inflammatory responses through NFκB inhibition: Implications for Crohn’s disease. Gut. 2000;47(3):397–403.

44. Elhenawy W, Tsai CN, Coombes BK, Elhenawy W, Tsai CN, Coombes BK. Host-Specific Adaptive Diversification of Crohn’s Escherichia coli Article Host-Specific Adaptive Diversification of Crohn’s Disease-Associated Adherent-Invasive Escherichia coli. Cell Host Microbe. 2019;25(2):301–312.e5.

45. Elhenawy W, Hordienko S, Gould S, Oberc AM, Tsai CN, Hubbard TP, et al. High-throughput fitness screening and transcriptomics identify a role for a type IV secretion system in the pathogenesis of Crohnâ€^™^s disease-associated Escherichia coli. Nat Commun. (2021):12(2032). doi:10.1038/s41467-021-22306-w

46. Boudeau Ã, Barnich N, Intestinale Â, De L, Pharmacie Â De. Type 1 pili-mediated adherence of Escherichia coli strain LF82 isolated from Crohn’s disease is involved in bacterial invasion of intestinal epithelial cells. Mol Micro. 2001;39(5):1272–1284.

47. Barnich N, Colombel J, Barnich N, Carvalho FA, Glasser A, Darcha C, et al. CEACAM6 acts as a receptor for adherent-invasive E. coli, supporting ileal mucosa colonization in Crohn disease. J Clin Invest. 2007;117(6):1566–1574.

48. Tropini C, Earle KA, Huang KC, Sonnenburg JL. The Gut Microbiome: Connecting Spatial Organization to Function. Cell Host Microbe. 2017;21(4):433–42. doi:10.1016/j.chom.2017.03.010

49. Human Microbiome Project Consortium. Structure, function and diversity of the healthy human microbiome. Nature. 2012;486:207–214. doi:10.1038/nature11234.

50. Lloyd-Price J, Mahurkar A, Rahnavard G, Crabtree J, Orvis J, Hall AB, et al. Strains, functions and dynamics in the expanded human microbiome project. Nature. 2017;550:61–66. doi:10.1038/nature23889.

